# Reinforcement Learning Identifies Age-Related Balance Strategy Shifts

**DOI:** 10.1101/2025.06.05.656344

**Authors:** Huiyi Wang, Jozsef Kovecses, Guillaume Durandau

## Abstract

Falls are one of the leading causes of non-disease death and injury in the elderly, partly due to the loss of muscle mass in a musculoskeletal disorder named sarcopenia. Studying the impact of this muscle weakness on standing balance through direct human experimentation poses ethical dilemmas, involves high costs, and fails to fully capture the internal dynamics of the muscle. To address these limitations, we employ neuromusculoskeletal modeling to explore the impact of sarcopenia on balance. In this study, we introduce a novel full-body musculoskeletal model comprising both the torso and lower limbs, with 290 muscle actuators controlling 23 degrees of freedom and supporting varying levels of sarcopenia. Using reinforcement learning coupled with curriculum learning and muscle synergy representations, we trained an agent to perform standing balance on a backward-sliding plate and compared its behavior to human experiments. Our results demonstrate that, without pre-recorded experimental data, both healthy and sarcopenic agents can reproduce ankle and hip balancing strategies consistent with experimental findings. Furthermore, we show that as the degree of sarcopenia increases, the agent adapts its balancing strategy based on the platform’s acceleration.

## I. Introduction

FALLINGS are one of the leading causes of non-disease death for individuals over the age of 65 years old [1]. Maintaining balance is a necessary component of achieving prolonged healthy living and increasing the quality of life. Failure to maintain balance can result in injuries such as bone fracture, traumatic brain injury, or related death [2]–[4]. Additionally, falls are responsible for 85% of injury-related hospital admissions among seniors, 95% of all hip fractures, and lead to over one-third of elderly individuals being placed in long-term care after hospitalization [5], [6]. One of the major accelerators of falls among the elderly is a progressive age-related, involuntary, and systemic loss of skeletal muscle mass and strength, also known as sarcopenia [7]. According to the Global Leadership Initiative in Sarcopenia, sarcopenia increases the risk of falls, fractures, and even mortality [8]– [10]. In order to protect the elderly population and promote healthy living, a proper understanding of balance strategies adapted to the decline of muscular strength is needed.

To maintain proper balance during standing, the body needs to ensure that its Center of Mass (CoM) stays within the Base of Support (BoS), the contact area between the feet and the ground. When standing, significant deviations from the center of the BoS lead to muscle activation around the hips (hip strategy), while minor disturbances are managed by the shank muscles that control the ankles (ankle strategy) [11], [12]. If balance cannot be regained from static balance, a stepping response is triggered (stepping strategy). Research indicates that as individuals age, their ability to quickly generate joint torques diminishes compared to younger adults due to neural or musculoskeletal disorders, which complicates the recovery of balance once a loss of balance begins [13], [14]. Therefore, a deeper understanding of how the central nervous system (CNS) coordinates balance in both young and elderly individuals is crucial. This knowledge could enhance clinical treatments for movement disorders and inform the development of controller designs for assistive technologies like exoskeletons and prostheses, which aim to counteract these disorders. Nevertheless, the specific mechanisms that adapt reactive motor behaviors in response to external disturbances remain poorly understood.

Balance has been traditionally studied by directly experimenting with human subjects. Using either a camera or a Motion Capture (MoCap) system and ground reaction plate, researchers are able to acquire the joint angles and torques through the use of inverse kinematics and inverse dynamics [15]–[17]. For example, [18] studies the effect of aging on stepping velocity to regain balance by recruiting both young and older subjects. [19] investigated the interchange between hip and ankle balancing responses on a backward translation plate. Moreover, study sometimes uses electromyography (EMG) recording to complement the joint measurement by providing information on the level of activation of the target muscles [19]. However, ethical concerns persist regarding the recruitment and experimentation of frail elderly individuals, and the general protocol for conducting human experiments is time-consuming and resource-intensive. In addition, the use of MoCap suffers from marker movements that produce inaccurate readings, and the measurement of deeper-level muscles requires invasive techniques.

In simulation, researchers commonly use a torque-driven single or double-linked inverted pendulum moving in the sagittal plane and parametrized using system identification methods to predict balancing responses without regard for muscle coordination [11], [15], [20], [21]. More recently, three-dimensional musculoskeletal models have been developed that reflect more accurately the neuromotor control and muscle-driven properties of the human body. For example, [22] and [23] have made use of the lower limb model in OpenSim [24] with 8 degrees of freedom (DoF) with 22 muscles to understand human balance in static and walking tasks. [25] and [26] studied a model with 23 DoFs and 92 Hill-type musculotendon actuators to understand the hip and ankle responses for reactive balance using forward dynamic simulations. Nevertheless, these models often feature limited trunk musculature which fails to fully capture the dynamics of torso movements.

In an overactuated system such as the human musculoskeletal system, multiple possible muscle activation patterns can achieve the same movement. Static or dynamic optimization techniques are traditionally used to identify muscle activation patterns based on a predefined cost function [25]–[27]. However, the design of those functions requires domain expertise and does not apply to functionally suboptimal behavior [28], [29]. Moreover, as the number of actuator combinations exponentially exploded with the increased number of muscle actuators, optimization methods failed to converge [30]. Reinforcement learning (RL) has been shown as a successful controller to tackle synthesizing physiologically feasible motion in high DoF biomechanical systems within a dynamics simulation environment (e.g., OpenSim, MyoSuite) [31], [32]. These simulation platforms have been used to reproduce human motion, leading to robust bipedal agility, including terrain running and direction changing [33]; and human postural control, such as balancing with robotic devices [34]. Compared to the dynamical optimization method, RL offers greater flexibility as it can perform predictive simulation and achieve generalization.

This paper presents a full-body musculoskeletal model consisting of 290 muscle actuators in the simulation platform MyoSuite [35]. We use this model to explore how the CNS coordinates the motor neurons to achieve standing balance with perturbation with different kinematic response strategies. We use RL with curriculum learning (CL) and muscle synergy representation (MSR) to generate a policy for balancing control. Our primary objective was to demonstrate RL can be a successful controller for an overactuated system like the human body for reactive balance responses in a three-dimensional muscle-driven model. Our secondary objective was to examine how these balancing responses change when the virtual agent experiences a reduction in maximal isometric force, simulating conditions such as sarcopenia. We hypothesized that our elderly musculoskeletal agent, even when simulating only muscle weakness, would exhibit distinct balancing behaviors that healthy agents. Further, we posited that RL could identify varying balancing strategies influenced by aging and perturbation intensity, rather than displaying uniform, monotonic responses. Our results demonstrate that without the help of any experimental data, our virtual agent produces the same non-stepping posture response as that of a human on a backward standing plate balancing task. Moreover, as the level of muscle force decreases, a continuous switch to hip strategy over ankle strategy is observed similar to what we observed in real human experiments [36].

## II. Methodology

### A. Musculoskeletal Modelling

Here, we present the first 3D full-body musculoskeletal model developed in MyoSuite. The model consists of both the torso model (MyoBack) [37] and the lower limb model (MyoLeg) [32]. The MyoBack consists of 210 muscle-tendon units (MTUs) 3 independent DoFs: flexion-extension, lateral bending, and axial rotation, which map to 21 DoFs. The model features a rigid pelvis and sacrum, five lumbar vertebrae, and a rigid torso consisting of a lumped thoracic spine and ribcage [38]. The MyoLeg contains 20 DoFs and 80 muscle-tendon units and was converted from the OpenSim full body model [39] via MyoConverter [40]. It consists of the right and left femur, patella, tibia/fibula, talus, calcaneus, and toes. Both models are driven by Hill-type musculotendon actuators following MuJoCo with inelastic tendons and no pennation angle. The sarcopenia condition is modeled by reducing the maximum isometric force of all muscle groups by *α ∈* (0, 1):

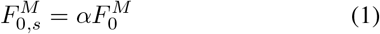

To reduce the dimension of the actuator for faster and better convergence, we utilize muscle grouping in the upper torso such that the muscles in the same group have the exact sample activation, as detailed in Figure 1 and Figure 2-A. We allow the left and right muscles of each group to be activated separately, giving in total of 24 muscle groups to control. In our simulation study, we consider four different models, where one serves as a reference without any agerelated modifications. The remaining three models simulate varying degrees of sarcopenia, characterized by decreases in the maximum isometric force each muscle can produce by 80%, 60%, and 40%, respectively. Other potential age-related factors, including sensorimotor delays, maximum contraction velocity, deactivation time constant, and tendon strain, are held constant across all models.

**Fig. 1.**
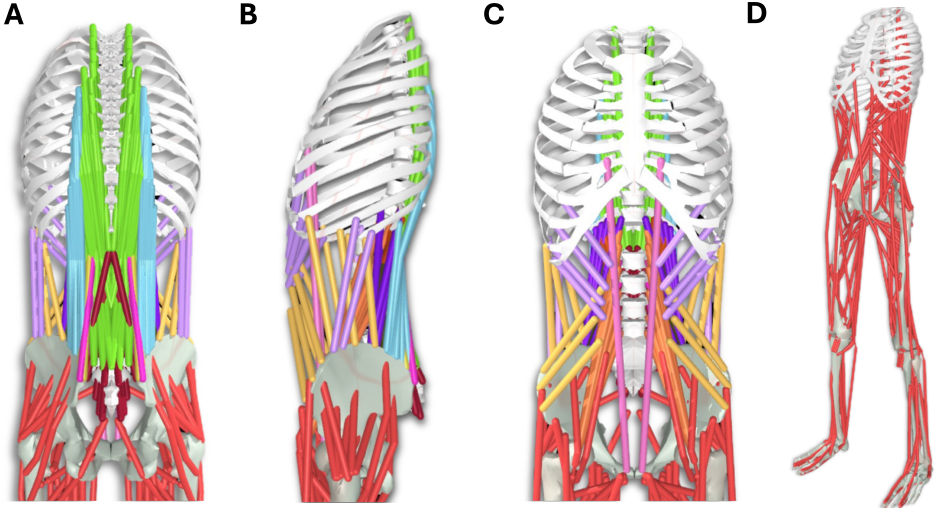
Full body musculoskeletal simulation model in MyoSuite presented in (A) back view (B) side view (C) front view (D) far view. The back muscles in (A) - (C) are color-coded according to the grouping. The lower back consists of seven main muscle groups: erector spinae (ES), rectus abdominis (RA), internal obliques (IO), external obliques (EO), psoas major (PM), quadratus lumborum (QL), and multifidus (MF). The ES group is further divided into four main components: the longissimus thoracis par lumborum (LTpL), the longissimus thoracis pars thoracis (LTpT), the iliocostalis lumborum pars thoracis (ILpT), and the iliocostalis lumborum pars lumborum (ILpL). Lastly, the QL group is divided into three subgroups: anterior fibers, lumbocostals, and the posterior fibers.

**Fig. 2.**
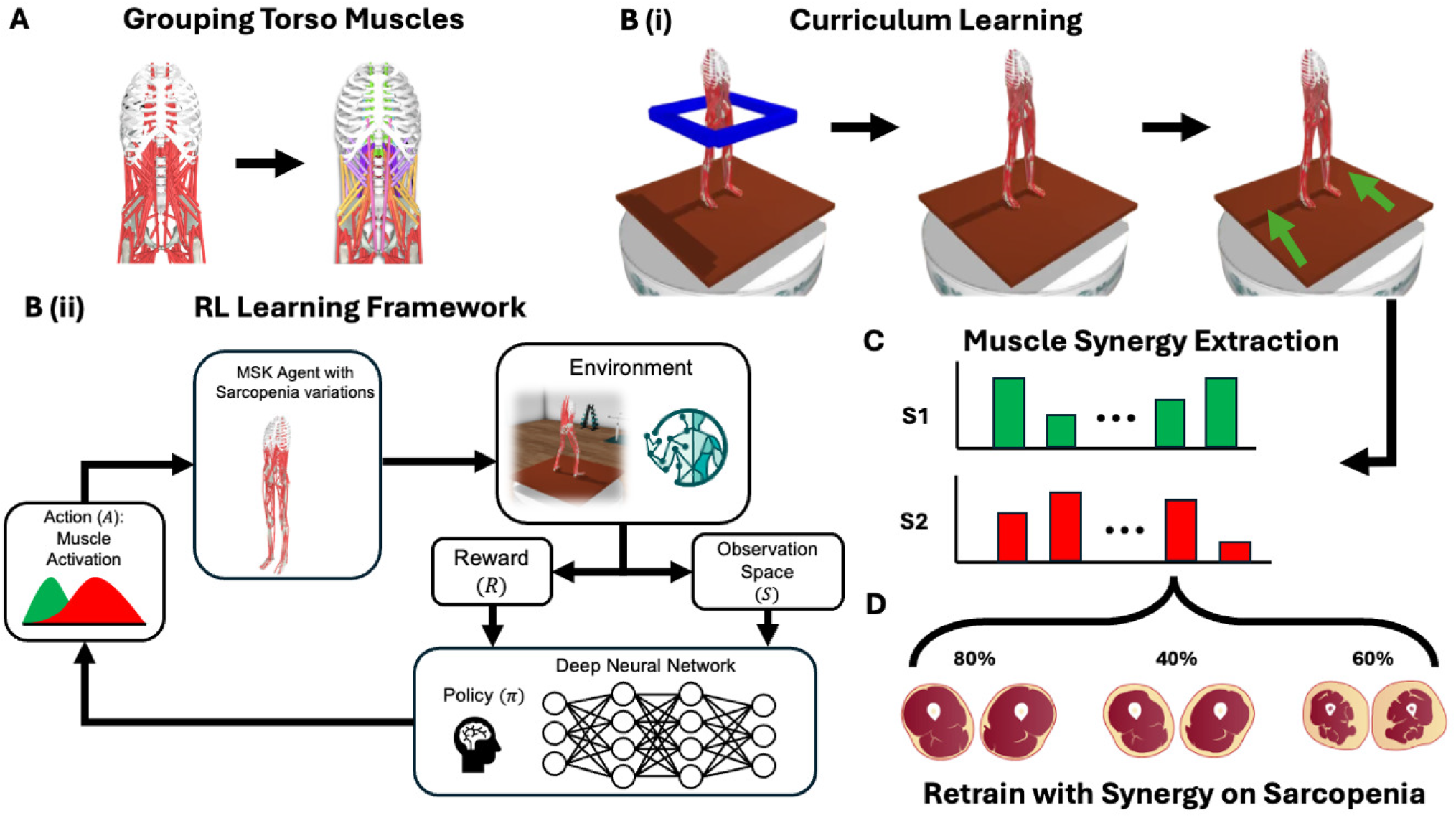
**A**: The 210 muscles in the torso are reduced into 24 groups via muscle grouping. **B-i**: Curriculum Learning based training is proposed to gradually maintain standing balance. **B-ii**: The deep reinforcement learning framework for a Musculoskeletal agent embedded within the MyoSuite simulation for healthy and sarcopenia agents when balancing on a backward translation plate. **C**: Extraction of muscle synergy pattern from healthy balance. **D**: Retraining the sarcopenia agent using the extract SAR representation.

### B. Reinforcement Learning

Reinforcement Learning (RL) is a machine learning method that allows an agent to learn through trial and error in an interactive setting, using feedback derived from its actions and experiences [41]. RL agents act within a Markov Decision Process (MDP), defined as the tuple (𝒮, 𝒜, 𝒫 ℛ,), where *s* ∈ 𝒮 is the observation state of the musculoskeletal agent including parameters such as joint positions, joint velocities, and metabolic costs, *a* ∈ 𝒜 is the different combination of the total muscle actuator activations presented in the MSK model, and the state-transition function 𝒫 = *P* (*s*_*t*+1_, *r*_*t*_ | *s*_*t*_, *a*_*t*_) represents the probability of moving from one state to another given a specific action. Without prior knowledge of 𝒫, the RL agent’s goal is to use the experience to maximize the expected rewards, 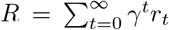, where *γ* ∈ [0, 1) is the discount factor. The goal of the RL agent is to learn a continuous control policy *π*^*\**^(*a* | *s*) that chooses actions that maximize the summed reward over time.

1. *Curriculum Learning:* Humans and animals tend to learn more effectively when examples are presented in a structured sequence that progressively introduces new concepts and increases in complexity, a method known as “curriculum learning” [42]. Similarly, machine learning benefits from a similar training strategy by starting with easier subtasks and gradually increasing the level of difficulty [43], [44]. In reinforcement learning, those types of methods are shown to provide better generalization and faster convergence. Formally, curriculum learning occurs when the entropy of the training distribution increases (i.e., additional training terms are introduced), and the weights of the new distribution monotonically increase as new curricula phases are introduced. The training of the full-body model was structured using a heuristically designed curriculum learning approach, segmented into three sequential stages: quiet balancing with external aid on the upper body, quiet balancing without external aid, and standing with perturbations, as shown in Figure 2-B. The policy’s training utilized reward functions based on the Euclidean distance between the agent’s target and current poses, modulated by curriculum-specific weights, *σ*_*i*_, as detailed in the following equation:

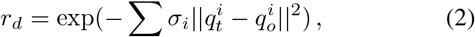

where 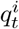 and 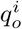 represent the target and observed joint positions, respectively. Additionally, a metabolic cost term was introduced to encourage energy-efficient movement.

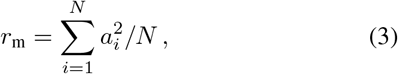

where *a*_*i*_ is the level of activation of the specific muscle and *N* is the total number of actuators presented in the model.
2. *Muscle Synergy Representation (MSR):* The overactuated nature of human muscles enables diverse movements with complex, distinctive activation patterns. However, analysis of EMG data shows that these patterns are often approximated by a linear combination of a few basic waveforms, indicating that the CNS efficiently resolves motor redundancy through a low-dimensional structure, a concept known as synergy. A prevalent method for identifying these underlying synergies involves applying Independent Component Analysis (ICA) to a subspace defined by Principal Component Analysis (PCA) [45], [46]. ICA is a technique for extracting features where the objective is to identify a linear representation of non-Gaussian data in such a way that the resulting components are statistically independent, or as close to independent as feasible [47]. Whereas PCA used to reduce the dimensionality of datasets, enhancing interpretability while simultaneously minimizing the loss of information [48].

In our study, we extracted the Muscle Synergy Representation (MSR) from a balanced, healthy agent trained using CL on a backward sliding plate, as shown in Figure 2-C. We selected the first 35 basis factors that account for 60% of the variance in the muscle stimulation of the simulated agent, which was tested to show the best balance between convergence speed and total return. This reduced representation was then used to retrain the healthy agent directly with perturbations, resulting in a more natural swing in body movements. In addition, the MSR, when applied to the sarcopenia models, achieves convergence with similar speed and return, whereas previously the CL was only able to converge to a sub-optimal local maximum (Figure 2-D).

### C. Simulation Setup

The model is placed on top of a backward support surface with the size of 80 *×* 80 *×* 3 (cm) that can only slide in the anterior-posterior (AP) direction. This support surface will undergo an anterior/posterior backward acceleration that is randomly sampled between the first 1 − 2 (*s*), with a perturbation acceleration randomly sampled between 1 − 30 (*m/s*^2^). The goal of the agent is to maintain the height of the CoM above its hip level for 5 seconds after the applied perturbation.

In our current task, the model was trained with an off-policy algorithm Soft Actor-Critic [49] using <monospace>Stable-Baselines3 </monospace>library [50]. The policy and Q-function use two fully connected hidden layers of 400 and 300 neurons respectively, with a linearly decreasing learning rate (*τ*) of 0.0005 and discount factor (*γ*) of 0.99. A ReLU activation function is used. Within the RL environment, the action space consists of a 35-dimensional continuous vector in the range, *a ∈* [− 1, 1]. The muscle actions are normalized to the range [0, 1] using:

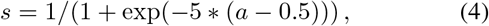

where *a* is the normalized action and *s* represents the muscle stimulation, with 0 and 1 indicating no stimulation and full stimulation, respectively. The policy has access to a 117-dimensional observation space, extensively describing the body’s joint position, joint velocity, the position of the target pelvis, and its distance from the current position.

The simulation results are validated against the experimental data collected with [26] and [51], which mirrors the setting of our simulation setup. In this study, subjects (young: 21 *±* 2 yrs; old: 67 *±* 3 yrs) were instructed to maintain balance on a platform that could randomly move in mediolateral and anterior-posterior directions with three acceleration profiles without stepping. In our simulation, we cross-match perturbation intensities by categorizing them as ‘low’ for acceleration ranges between 0 − 10*m/s*^2^ and ‘high’ for those between 20 − 30*m/s*^2^. The three elderly models are compared together with the older subject data since the muscle level of human subjects is not measured.

## III. Results

Figure 3 shows the ankle and hip joint angle responses of a healthy agent across two perturbation levels. Under low perturbation conditions, joint angles remain stable across different accelerations 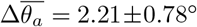 and 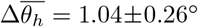 for anterior perturbation), while joint moments correlate positively with increased acceleration. In high perturbation scenarios, both joint angle and joint torque directly correlate with increasing acceleration. Table I summarizes the comparison of the averaged joint range between the experimental and simulation data for the healthy agent, with the majority of the data matching within one standard deviation.

**TABLE I.**
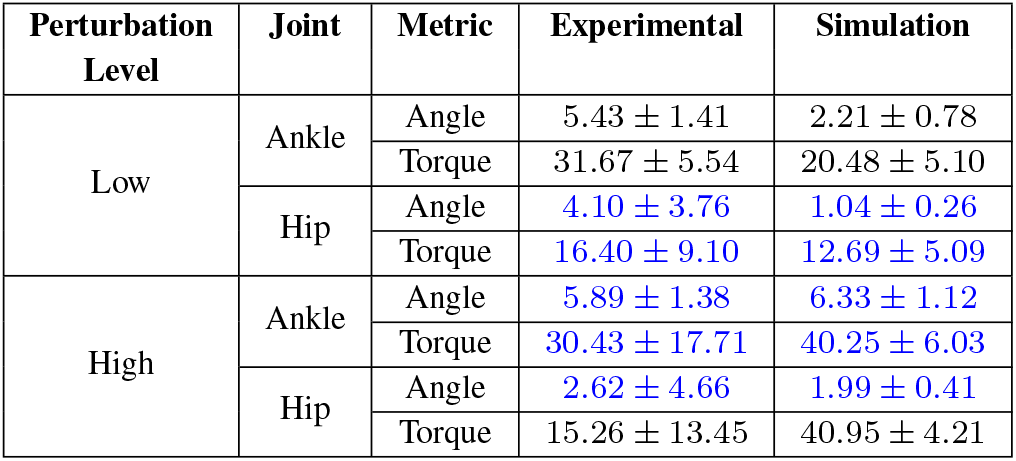
Comparison of averaged range of ankle and hip joint angles 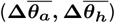 and torques 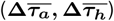 between experimental values and simulation in healthy under low and high anterior perturbations. Data highlighted in blue indicates where simulation and experimental data match within one standard deviation. Angles are measured in degrees (°) and torques in Newton-meters (Nm).

| Perturbation Level | Joint | Metric | Experimental | Simulation |
| --- | --- | --- | --- | --- |
| Low | Ankle | Angle | $5.43 \pm 1.41$ | $2.21 \pm 0.78$ |
| | | Torque | $31.67 \pm 5.54$ | $20.48 \pm 5.10$ |
| | Hip | Angle | $4.10 \pm 3.76$ | $1.04 \pm 0.26$ |
| | | Torque | $16.40 \pm 9.10$ | $12.69 \pm 5.09$ |
| High | Ankle | Angle | $5.89 \pm 1.38$ | $6.33 \pm 1.12$ |
| | | Torque | $30.43 \pm 17.71$ | $40.25 \pm 6.03$ |
| | Hip | Angle | $2.62 \pm 4.66$ | $1.99 \pm 0.41$ |
| | | Torque | $15.26 \pm 13.45$ | $40.95 \pm 4.21$ |

**Fig. 3.**
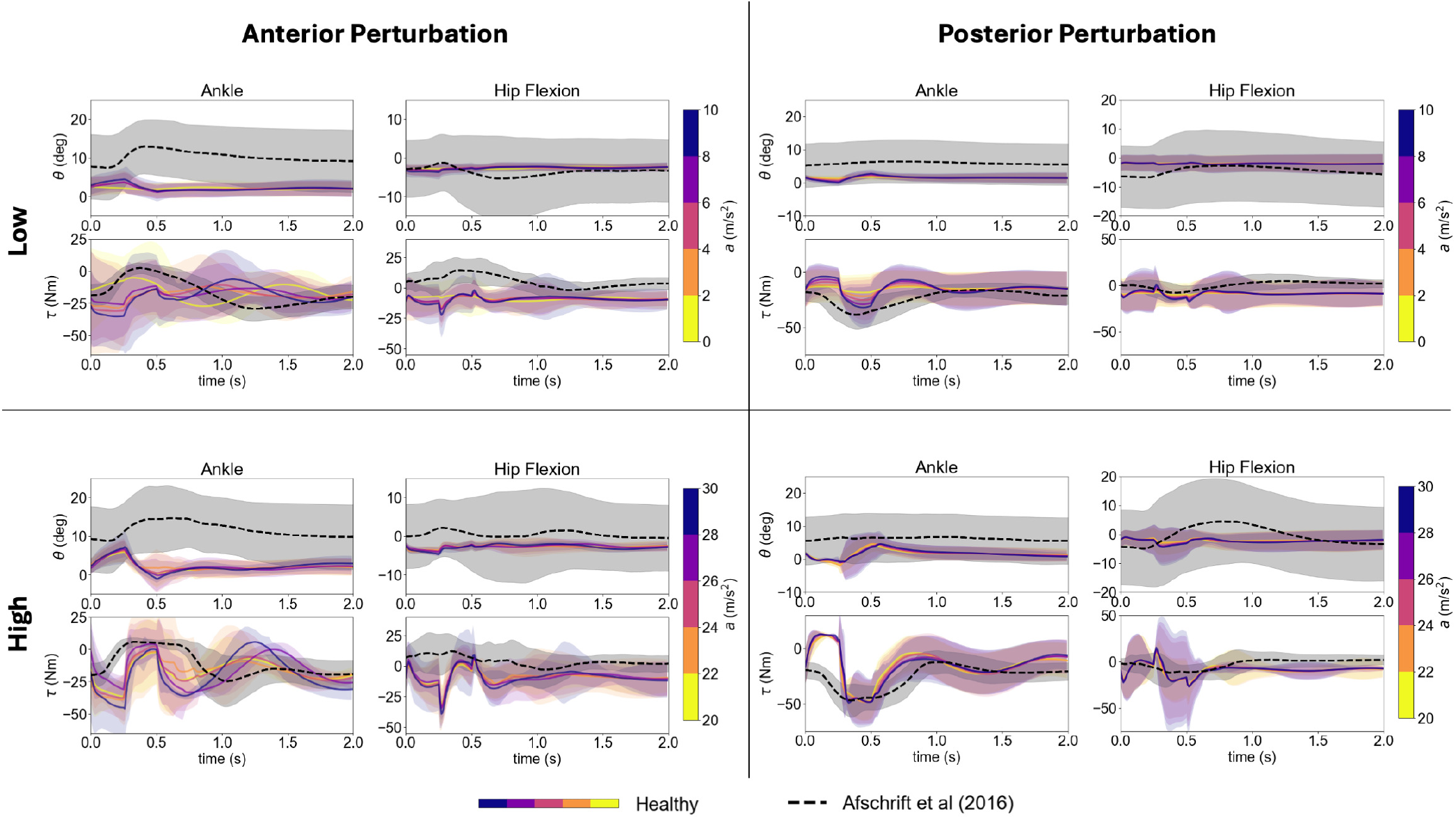
Joint information for healthy agents when experiencing an anterior (left) or posterior (right) perturbation. Line colors correspond to the colorbar scale shown to the right of each plot. The upper row shows the joint kinematics under a low perturbation (0 - 10 ***m/s***^***2***^) while the lower row shows the joint kinematics for a high perturbation (20 - 30 ***m/s***^***2***^). The colored solid line represents the mean joint angle across 50 trials, whereas the shaded area represents mean ***±*** 1 standard deviations (std). The dashed black lines are experimental data from [26], with the shaded grey area to be ***±*** 1 std. Data are aligned such that the onset of perturbations is marked at 0 seconds.

For sarcopenia agents, as shown in Figure 4, the averaged ankle joint value deviates from the experimental data by around 10^*°*^, indicating a more upright position in the simulation. For low perturbations, sarcopenic agents maintain stable joint angles and torques (e.g., 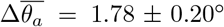 and 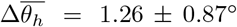) in contrast to the large joint angle and torque in experiment (Table II, upper row); whereas for high perturbations, significant peaks occur in both ankle and hip joints (e.g., 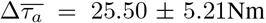 and 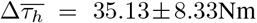), aligning more closely with the experimental trend.

**TABLE II.** Comparison of averaged range of ankle and hip joint angles 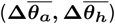 and torques 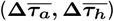 between experimental values and simulation for sarcopenia models under low and high perturbations. The three sarcopenia models are treated as one. Data highlighted in blue indicates where simulation and experimental data match within one standard deviation. Angles are measured in degrees (°) and torques in Newton-meters (Nm).

| Perturbation Level | Joint | Metric | Experimental | Simulation |
| --- | --- | --- | --- | --- |
| Low | Ankle | Angle | $6.34 \pm 1.49$ | $1.78 \pm 0.20$ |
| | | Torque | $39.92 \pm 8.72$ | $8.83 \pm 1.53$ |
|  | Hip | Angle | **$3.96 \pm 1.80$** | **$1.26 \pm 0.87$** |
| | | Torque | $25.07 \pm 4.48$ | $8.05 \pm 2.30$ |
| High | Ankle | Angle | $10.27 \pm 1.05$ | $6.80 \pm 0.49$ |
| | | Torque | $42.47 \pm 7.42$ | $25.50 \pm 5.21$ |
|  | Hip | Angle | **$4.99 \pm 0.58$** | **$4.52 \pm 2.41$** |
|  |  | Torque | **$23.08 \pm 10.16$** | **$35.13 \pm 8.33$** |

**Fig. 4.**
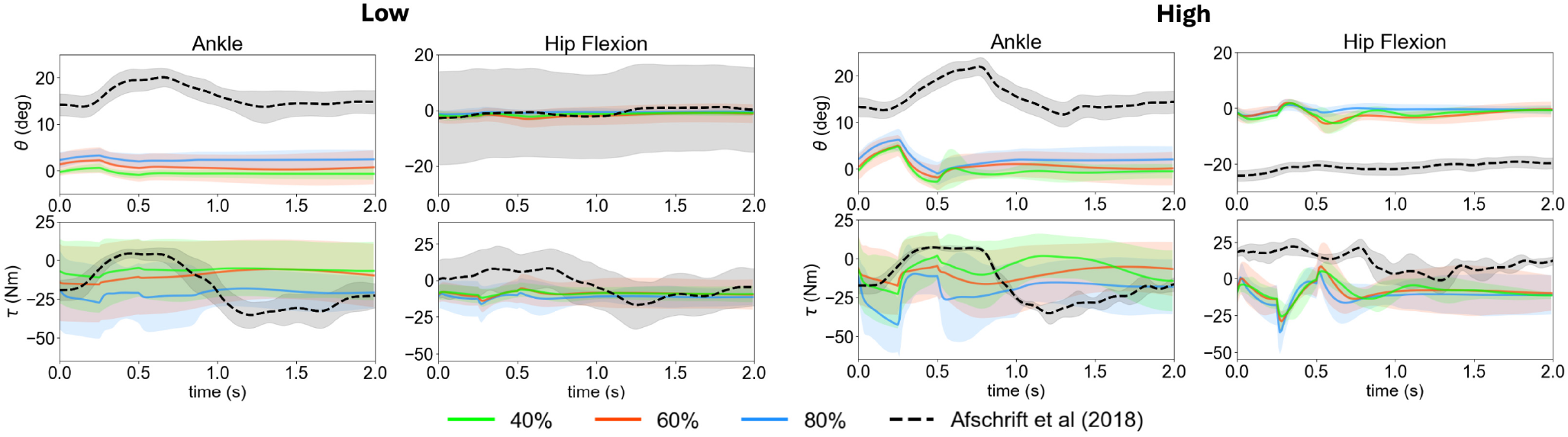
Joint information for sarcopenia agent with different muscle mass percentage in comparison to human experimental data collected [51] facing an anterior perturbation. The solid lines are mean values of joint information across high (right) and low (left) perturbation levels for ankle and hip strategy respectively, with the shaded area to be ***±*** 1 std.

The simulation results of trunk flexion-extension is shown in Figure 5-A. While the average trunk flexion angle in the sarcopenia model aligns with experimental data of 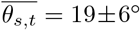, the healthy model leans forward more than observed in healthy human subjects: 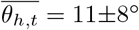,indicating a more conservative, energy-inefficient stance likely due to perturbation anticipation^1^. Additionally, all models experience a slight increase in trunk flexion range as the perturbation level increases, with the healthy agent showing the least variation. In terms of the Euclidean displacement of the center of mass (CoM), as shown in Figure 5-B, the healthy, 80%, and 60% muscle force models maintain stability by staying within 1 cm of the starting position. A slight increase in the center quartile is observed in these models as the level of perturbation intensifies. However, the range of CoM of the 40% muscle force model reaches more than 10cm when facing a perturbation level between 30 − 40*m/s*^2^, resulting in a fall.

**Fig. 5.**
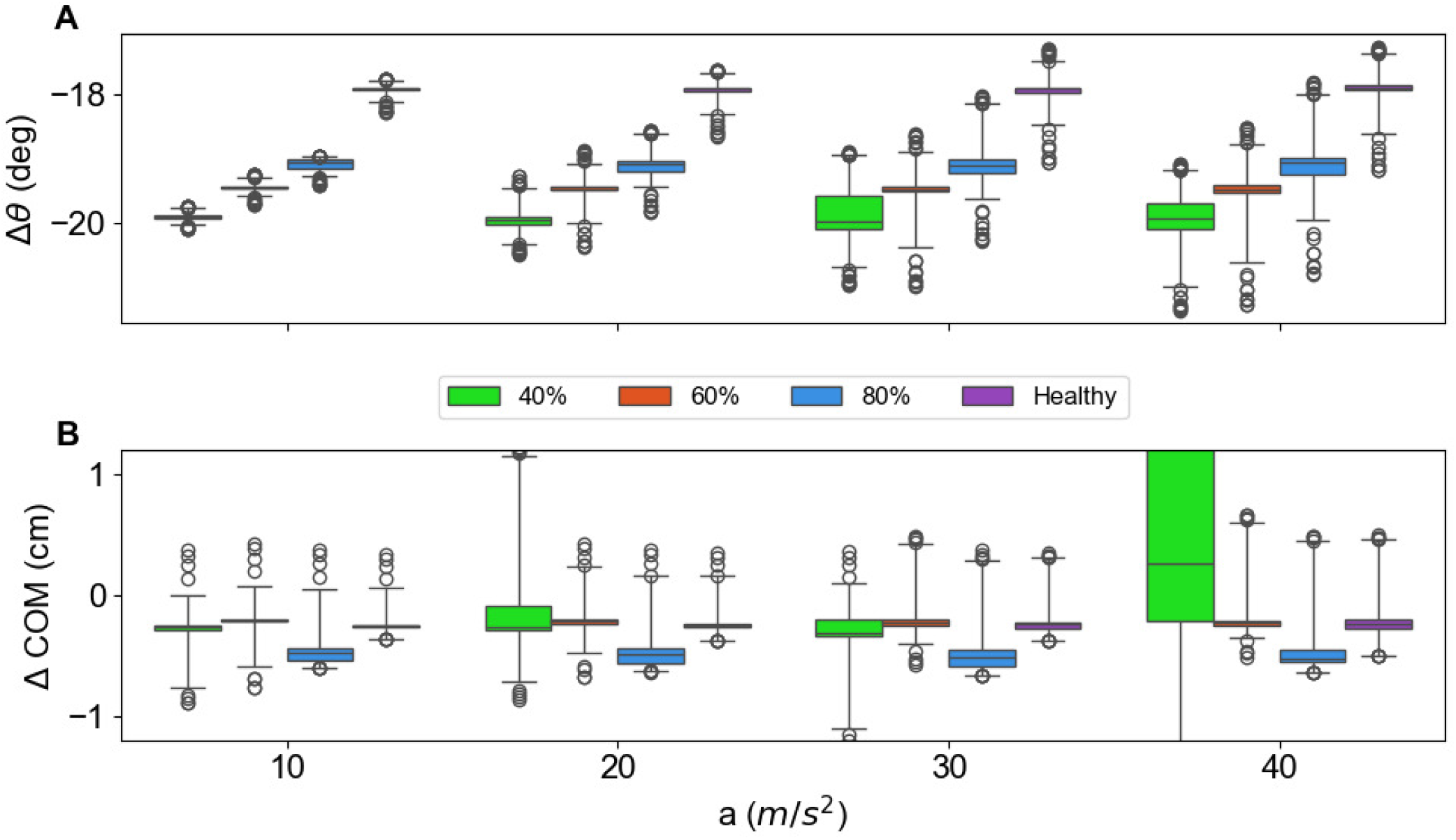
The range of (A) trunk flexion-extension angle and (B) the Euclidean displacement of COM as a function of the platform acceleration for agents with four different muscle force levels. Four trials with **40%** muscle force level experienced falling. The graph is truncated to enhance visibility of key data points; additional data extending beyond the x-axis range is omitted for clarity.

Figure 6 indicates the maximum range of hip and ankle angles experienced by the agent in the face of perturbations. We define the onset of the hip strategy when the agent’s hip flexion exceeds 4° during balance tasks. A healthy agent can withstand perturbations up to 35 *m/s*^2^ using only ankle strategies, while sarcopenic agents resort to hip strategies at lower thresholds of 25 *m/s*^2^, 15 *m/s*^2^, and 0 *m/s*^2^, corresponding to 80%, 60%, and 40% of muscle force, respectively.

**Fig. 6.**
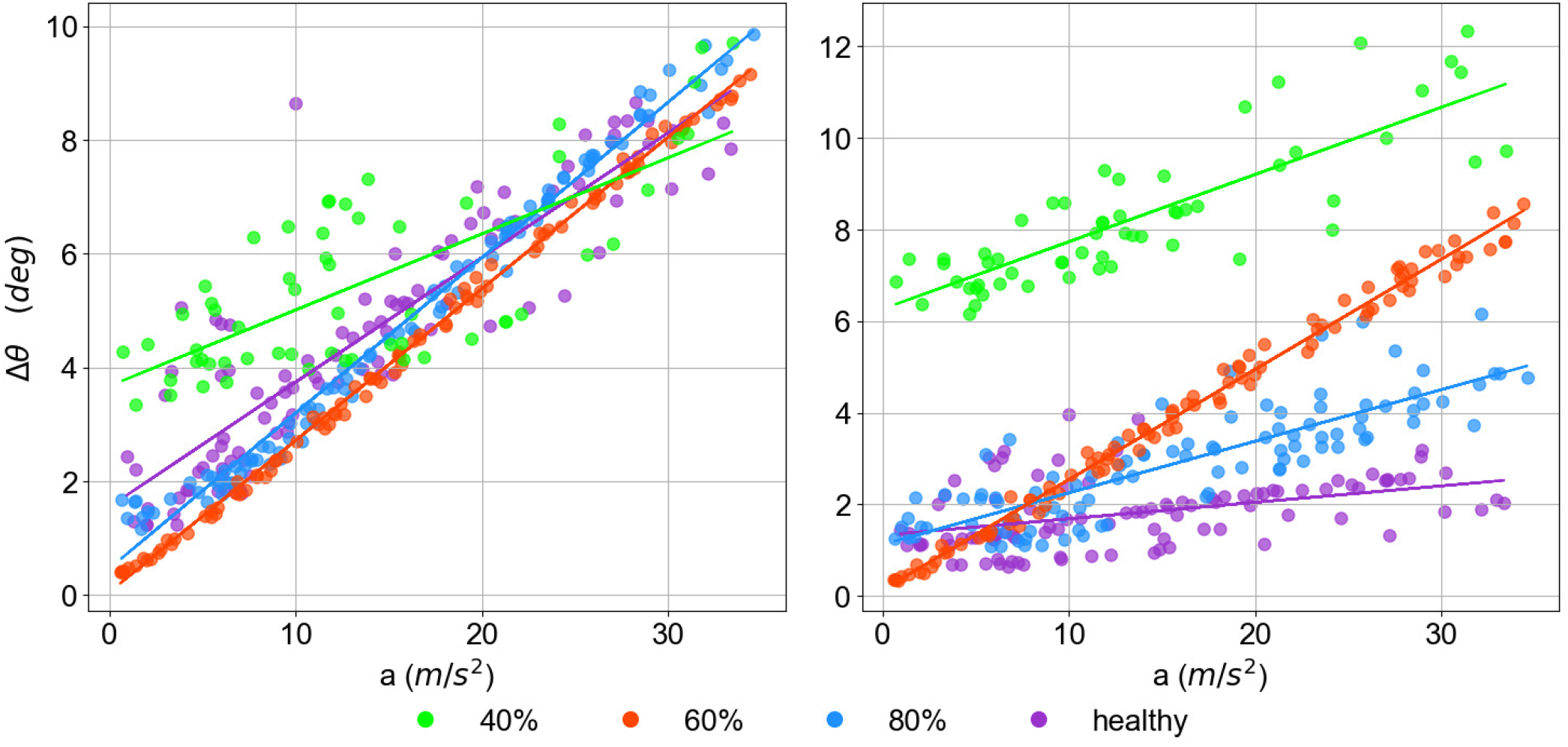
The maximum displacement in joint angle for ankle (left) and hip flexion (right) as a function of perturbation acceleration. A comparison between the level of sarcopenia (e.g., 80 corresponds to 80% of total maximal isometric force), and the maximum displacement of joint torque and moment exerted. The colored line represents a linear fit to the scattered data.

## IV. Discussion

### A. Kinematics of the postural responses

We compare the kinematic response when facing anterior and posterior perturbations independently with experimental data of the healthy model. The virtual agent exhibits joint angles and joint torques that are opposite in direction between these two conditions, aligning with human subjects. Notably, minimal hip joint torque 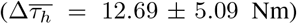 during low-level perturbations indicates a primary reliance on ankle strategies for balance. In contrast, during high-level perturbations, significant increases in hip angle and torque 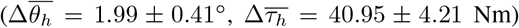 suggest a shift to hip-based balancing strategies. Additionally, the use of hip torque is always accompanied by ankle torque and is not observed individually.

While experimental data indicate that elderly subjects exhibit high hip torque in both low and high perturbation conditions, our simulations reveal a continuous increase in the hip torque exerted. This observation indicates that, in real experiments, elderly subjects prefer using hip strategies for balance even under low perturbations, where ankle strategies might be adequate. This preference could be due to a psychological fear of falling when using only ankle strategies. Additionally, compared to healthy individuals, elderly subjects exhibit a longer corrective response to platform translations. Our virtual agent does not replicate this trend, suggesting that the extended response time in elderly people may be related to sensorimotor delays not represented in our simulation.

In addition, our model demonstrates increased trunk flexion extension as the sarcopenia level increases. This is similar to the condition of hyperkyphosis observed in the elderly characterized by a forward-leaning posture that reduces mobility and increases the risk of falls due to weakness inspinal extensor muscles [52]. Moreover, the stability of CoM across the healthy, 80% and 60% force models suggested our model acquired robust policy in standing perturbations. In contrast, the 40% force model, which results in 4 cases of the falling, indicates that a severely reduced muscle force model contributes to the decreased ability of the model to control the whole body CoM. Notably, this model with the most severe sarcopenia employs the hip strategy even at minimal perturbations, indicating that the hip strategy is more capable of accelerating the CoM and stabilizing posture even with weaker muscle forces. The ankle angle displacement in the weakest force model reaches as high as 4^*°*^ at low perturbations, suggesting an early adoption of a combined ankle and hip strategy compared to the other models. These simulation results are consistent with the findings from other studies (e.g., [36], [53], [54]), which observed a shift from ankle to hip strategy for balance occurs at a lower perturbation in older than in young healthy adults.

### B. Evaluation of Hypotheses

The main goal of this study is to understand whether an RL controller applied to a model with more than 290 muscle actuators can accomplish a standing balance task on a backward support surface. We demonstrated that by integrating CL with MSR, meaningful synergistic representations can be extracted and reused on sarcopenia models. Our results show that both healthy and sarcopenic virtual agents, even with their maximum isometric force reduced by over half, could handle high perturbations without relying on experimental data. In line with our initial hypothesis, different sarcopenia agents exhibited distinct hip and ankle joint kinematics even when simulating muscle weakness as the sole age-related musculoskeletal disorder.

Our secondary goal is to show that our musculoskeletal agents can display a continuum of responses in the face of different perturbation magnitudes. While the virtual agent could have relied on a single strategy for balance, the inclusion of a metabolic cost reward term encourages it to adapt and optimize its responses by maintaining an upright posture while minimizing muscle activation. Our results indicate that the virtual agent can switch between hip and ankle strategies based on the perturbation intensity, aligning with the findings of [36]. The hip strategy is more prevalent in elderly subjects, even under low perturbation conditions, suggesting that our RL agent mirrors the behavior of CNS to prioritize hip strategies to maintain stability in the elderly.

### C. Algorithm Performance

Our model, based on only two simple reward terms: minimization of joint angles and total metabolic costs, generates a continuum of kinematic responses that closely resemble experimentally observed variations in combinations of hip and ankle strategies. This method offers a straightforward approach compared to earlier simulations (e.g., [25]), which defined a complex cost function with three components (CoM excursion, trunk orientation excursion, and muscle stress) and over ten variables to optimize. Similarly, [55] utilized a model with four distinct reward terms, including target posture, torque, body upright position, and extrapolated CoM. Our approach requires less domain expertise and achieves convergence with minimal sensitivity to hyperparameter adjustments. Additionally, our model demonstrates robustness against perturbations of varying durations and initial onset times that are not presented during training without specific fine-tuning.

The selection of the number of MSRs influences the performance of our agent in balance tasks. While varying the number of MSRs all result in a stable, quiet balance, fewer representations (*<* 30) lead to falling with high perturbations. This indicates that reflex behaviors in balancing require a more complex MSR than is necessary for simple quiet standing. Conversely, a higher number of MSRs (*>* 50) requires a long convergence time (up to 5 million timesteps), yet it does not significantly enhance performance in maintaining balance under forces up to 40*m/s*^2^. In addition, using MSR shows higher robustness in producing feasible balancing behavior for elderly models in comparison to CL. When training elderly models to balance from scratch using CL, they often converge to local minima, such as relying on external aid (blue bars shown in Figure 2-Bi), rather than learning to stand independently.

### D. Limitations and Future Outlook

Our study has several limitations. Primarily, our musculoskeletal model employs a generic representation rather than being scaled to the weight and size of individual participants. Previous research by [56] and [57] demonstrates that body mass significantly influences the extent of movements required to maintain postural balance, accounting for over 50% of stability variations among adults. Furthermore, [58] underscores the significance of gender differences in balance maintenance capabilities among the elderly. Personalizing the model to reflect these gender-specific differences could provide deeper insights into the interplay between balance ability and agerelated factors.

Secondly, our study does not incorporate specific musculoskeletal and neurological disorders like sensorimotor delays and reduced muscle contraction velocity in simulations of elderly models. Research such as [23] has shown that including these factors provides a more accurate representation of elderly fall data during walking than modeling based solely on age-related muscle force decline. Although results by [59] suggested that loss of muscle strength and mass is the main cause of age-related inefficiency and speed in walking, future iterations could benefit from integrating additional neurological diseases.

Lastly, although our study shows a high overlap in the kinematic responses between simulation and experimentation, the muscular activation patterns are physiologically unrealistic in comparison to the experimentally collected data. As the human MSK system is overactuated, multiple possible muscle activation patterns that can achieve the same kinematic responses, which our current RL reward design could not capture. Future work will focus on integrating EMG data directly into the training process, which could help align the simulated muscle activation patterns more closely with experimental observations and enhance our model’s potential for studying neuromusculoskeletal disorders and developing assistive technologies.

## V. Conclusion

In this study, we present a full-body musculoskeletal model with 290 muscle actuators in the simulation platform MyoSuite. We demonstrate that reinforcement learning, coupled with curriculum learning and muscle synergy representation, can be a successful controller for an overactuated system like the human body for reactive balance responses in a three-dimensional muscle-driven model. Our results show that without the help of any experimental data, our virtual agent produces the same non-stepping posture response as that of a human on a backward standing plate balancing task. Moreover, as the level of muscle force decreases, a continuous switch to hip strategy over ankle strategy is observed. Our study provides the foundation for future research on using reinforcement learning to understand human balance control and explore the influence of aging on balance maintenance.

## Footnotes

1 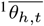 and 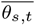 are reported in human experiment with high perturbations.

