## Supplementary figures and images for "Reinforcement Learning Identifies Age-Related Balance Strategy Shifts"

### Supplementary Movie

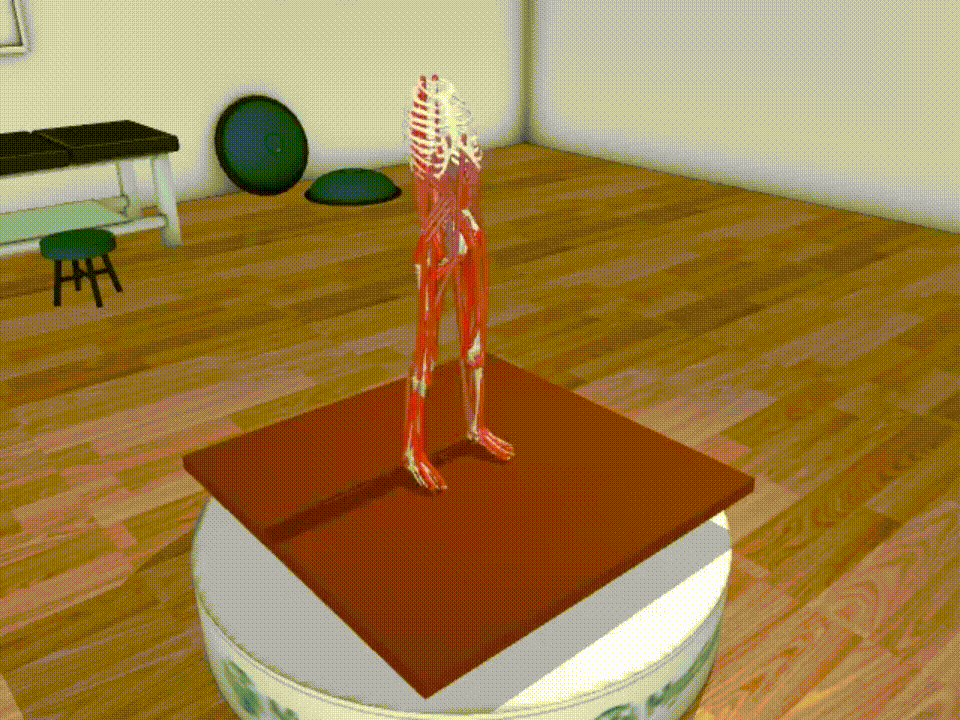
